# Dose-dependent spatiotemporal responses of mammalian cells to an alkylating agent

**DOI:** 10.1101/215764

**Authors:** Ann Rancourt, Sachiko Sato, Masahiko S. Satoh

**Affiliations:** Glycobiology and Bioimaging Laboratory of Research Center for Infectious Diseases; Laboratory of DNA Damage Responses and Bioimaging, CHUQ, Faculty of Medicine, Laval University. 2705 Boulevard Laurier, Quebec, Quebec, G1V 4G2, Canada

**Keywords:** Carcinogens, single-cell tracking, spatiotemporal responses

## Abstract

Cultured cell populations are composed of heterogeneous cells, and previous single-cell lineage tracking analysis of individual HeLa cells provided empirical evidence for significant heterogeneity of cell fates. Nevertheless, such cell lines have been used for investigations of cellular responses to various substances, resulting in incomplete characterizations. This problem caused by heterogeneity within cell lines could be overcome by analyzing the spatiotemporal responses of individual cells to a substance. However, no approach to investigate the responses using spatiotemporal data is currently available. Thus, the current study aimed to analyze the spatiotemporal responses of individual HeLa cells to cytotoxic, sub-cytotoxic, and non-cytotoxic doses of the well-characterized carcinogen, *N*-methyl-*N*'-nitro-*N*-nitrosoguanidine (MNNG). Although cytotoxic doses of MNNG are known to induce cell death, the single-cell tracking approach revealed that cell death occurred following at least four different cellular events, suggesting that cell death is induced via multiple processes. We also found that HeLa cells exposed to sub-cytotoxic doses of MNNG were in a state of equilibrium between cell proliferation and cell death, with cell death again induced through different processes. However, exposure of cells to non-cytotoxic doses of MNNG promoted growth by reducing the cell doubling time, thus promoting the growth of a sub-population of cells previously recognized as putative cancer stem cells. These results demonstrate that the responses of cells to MNNG can be analyzed precisely using spatiotemporal data, regardless of the presence of heterogeneity among cultured cells, suggesting that single-cell lineage tracking analysis can be used as a novel and accurate analytical method to investigate cellular responses to various substances.

## Introduction

Cellular responses to genotoxic insults have been investigated using various end-point analyses that measure alterations induced in cells at a specific moment in time, and then deduced the likely cellular responses by evaluating the data obtained at different time points. These results may be valid if all the individual cells within a cell population share similar characteristics and the cellular responses are induced in a stochastic manner. However, cell-to-cell heterogeneity has been demonstrated among cultured cells, though no empirical data revealing the spatiotemporal heterogeneity between individual cells has yet been available. We previously developed a novel chronological, single-cell lineage tracking analysis system that could record cellular events and movements of live cultured cells in a continuous manner using differential contrast imaging (DIC) (1). Resulting live cell imaging videos, which contained multidimensional information, including morphology of cells, position of individual cells within a cell population and types of cellular events occurred in a cell. This approach allowed us to extract critical information of individual cells by performing single-cell tracking and creating cell-lineage database. The cell population could then be characterized using this database. Using this system, we previously showed significant cell-to-cell heterogeneity in cultured HeLa cells and demonstrated that the fates of individual cells were diverse, indicating that the HeLa cell line comprises a highly heterogeneous population of cells (1). Moreover, some cells, tentatively referred to as putative cancer stem cells, had a reproductive ability >20 times higher than that of other cells (1). The A549 lung carcinoma cell line and mouse embryonic fibroblasts have also been shown to comprise heterogeneous cells (2, 3). These observations suggest that individual cells within a cell line may not share similar characteristics, and an accurate determination of cellular responses to genotoxic insults thus requires investigation of the spatiotemporal responses of individual cells to the insult. However, no such approach has yet been fully developed.

The alkylating agent *N*-methyl-*N*'-nitro-*N*-nitrosoguanidine (MNNG) is one of the best-characterized carcinogens, mutagens, and DNA-damaging agents. MNNG produces methylated DNA bases, such as *N*^7^-methyl guanine, which can be efficiently repaired by base excision repair (4–8). Exposure of cells to cytotoxic doses of MNNG activates poly(ADP-ribose) polymerase-1 during the repair process, and the synthesis of poly-ADP ribose polymer from NAD^+^ leads to cell death (CD) due to the depletion of intracellular NAD^+^ or activation of ADP-ribose polymer-induced apoptosis (9–14). Another MNNG-induced methylated DNA base, *O*^6^-methyl guanine, causes formation of *O*^6^-methyl G:T and *O*^6^-methyl G:C mismatches when the DNA containing *O*^6^-methyl G is replicated (15, 16). It has been suggested that these mismatches are repaired by mismatch correction (17–19), which has also been suggested to be involved in CD induced by low levels of alkylating agents (20–23). Clinically relevant doses of alkylating agents, as used for anticancer treatment, also induce this type of CD (24), although various models have been proposed regarding the induction of CD (20–23, 25–31).

In this study, we analyzed the spatiotemporal responses of HeLa cells to exposure to non-cytotoxic, sub-cytotoxic, and cytotoxic doses of MNNG using single-cell lineage tracking analysis, to develop an approach for the characterization of cellular responses in a heterogeneous cell population. Our results and analyses at single cell level suggest that individual cells respond to MNNG via different processes and their responses to different doses also significantly vary, while the lineage-based analyses reveal some levels of consistency in dose-dependent responses, which had not been previously revealed by classic population-based experiments and analyses. Given that classical end-point assays, which have previously been used to investigate various biological processes, are unable to determine the specific processes whereby cells eventually undergo CD, spatiotemporal data will be essential for conducting accurate investigations of cellular responses to various external substances.

## Results

### Accuracy of analysis using spatiotemporal data and effect of MNNG exposure on cell-doubling time

HeLa cells were exposed to three different doses of MNNG for 30 min, as follows: non-cytotoxic (1 μM dose at which cell proliferation was not reduced; MNNG1), sub-cytotoxic (2 μM dose at which CD was induced but proliferation also occurred; MNNG2), and cytotoxic dose (5 μM dose at which CD predominantly occurred; MNNG5) (see Movies S1–4, and Movie S5 for exposure to lethal dose of MNNG, 40 μM). We referred to cells identified at time point 1 as progenitor cells, and these cells and their progeny were then tracked over time to create a cell-lineage database and maps (Figs. S1–4). We previously confirmed the reproducibility of the single-cell lineage tracking analysis by evaluating cell-growth curves based on three independent imagings of untreated cells (1). We extended this analysis to characterize the responses of HeLa cells to MNNG exposure by simultaneous imagings of control and exposed cells. We evaluated cell doubling time in addition to classic cell growth curves, given that cell growth obtained by plotting the number of cells at multiple time points was influenced by the occurrences of multipolar cell division (MD, mainly tripolar cell division (32, 33)), CD, and cell fusion (CF). The mean cell doubling times determined from cell-tracking data of individual control and MNNG1 cells from two independent imagings are shown in Fig. 1A. The cell doubling time of MNNG1 cells was consistently shorter than that of the control cells (Fig. 1A, control vs. MNNG1). A comparison between individual control imagings showed a 1.124-fold difference in mean cell doubling times; however, because the statistical analysis involved >1,000 data points, a 1.124-fold difference between imagings could only be detected by single-cell lineage analysis. We confirmed that this degree of variation did not impact on the analysis of responses to MNNG1 by normalizing the cell doubling time of Imaging 2 of control and MNNG1 cells by a factor of 1.124 (Fig. 1A, Imaging 2, Normalized). There was no significant difference between Imaging 1 and normalized Imaging 2, confirming that variation between imagings did not affect the analysis of cellular responses to MNNG1. Furthermore, these results suggest that MNNG1 reduced the cell doubling time. We found similar reductions using merged cell lineage databases for Imaging 1, control and MNNG1 cells and their Imaging 2 counterparts (Fig. 1B), and we therefore used merged databases for subsequent analyses. Similar to the response to MNNG1, the mean cell doubling time of cells exposed to MNNG2 was significantly shorter than that of the control cells (Fig. 1B), while the cell doubling time of cells exposed to MNNG5 was prolonged (Fig. 1B). These observations were consistent with earlier reports that cytotoxic doses of MNNG perturbed progression of the cell cycle (34). We also analyzed cell doubling time using a Gaussian distribution (Fig. 1C-D) and confirmed that cells exposed to MMNG1 and MMNG2 showed shorter cell doubling times, suggesting that spatiotemporal data obtained by single-cell lineage tracking analysis allowed a precise analysis of the effect of exposure to MNNG on cell doubling time.

**Fig. 1.**
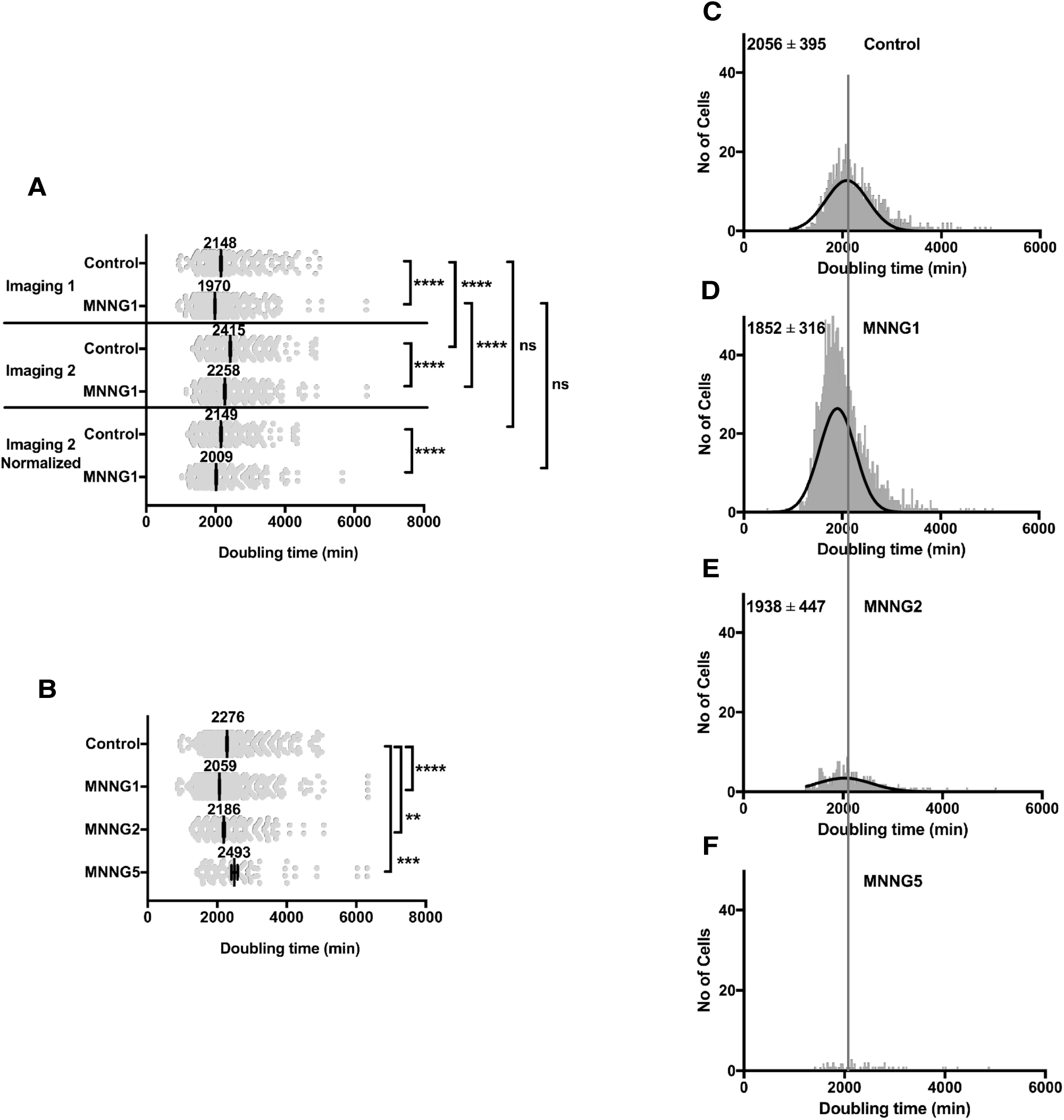
Analysis of cell-doubling time. **A**. Two sets of videos for Control and MNNG1-exposed cells were created by independent long-term live cell imaging and used for cell tracking analysis (Imagings 1 and 2). Cell doubling times analyzed by Imaging 2 were normalized by a factor of 1.124 (Imaging 2, Normalized). One-way ANOVA was performed; ns: not significant and ****p<0.0001. Results shown as mean ± standard error (SEM). **B**. A cell lineage database was created by merging the two cell lineage databases. Cell doubling times were analyzed for Control, MNNG1-, MNNG2-, and MNNG5-exposed cells. Student’s *t*-tests were performed; **p<0.01, ***p<0.001 and ****p<0.0001. Results shown as mean ± SEM. **C-F**. Cell-doubling time distributions of Control cells (**C**) and cells exposed to MNNG1 (**D**), MNNG2 (**E**), and MNNG5 (**F**) are shown. Mean cell doubling times and standard deviations were calculated (**C-E**) using the Gaussian distribution (Prism 7).

### Effect of MNNG exposure on rate of cell population expansion using spatiotemporal data

We also evaluated the effect of MNNG on the rate of cell population expansion using spatiotemporal data. Conventional methods determine the rate of cell population expansion by time course analysis, in which the number of cells at each time point was determined by averaging the results from multiple cell countings. In contrast, single-cell lineage tracking analysis determined the numbers of progenitor cells and/or progeny of each cell lineage at each time point (10 min) (see cell lineage maps for Figs. S1–4), and used the total for each time point to draw the cell population expansion curves (Fig. 2A). We first evaluated the rate of cell population expansion of MNNG1 cells and showed that MNNG1 treatment significantly increased the numbers of cells at 6,000 and 8,500 min (p <0.001) (Fig. 2B), suggesting that MNNG1 could promote cell proliferation. In contrast, MNNG2 treatment significantly suppressed cell proliferation (Fig. 2B). There was a slight increase in cell number at around 2,500 min (Fig. 2C) and the overall increase at 8,500 min was only 1.5-fold (Fig. 2A), implying that both cell proliferation and CD occurred simultaneously following exposure to MNNG2. However, cytotoxic doses of MNNG (Fig. 2A, MNNG5) inhibited cell population expansion.

**Fig. 2.**
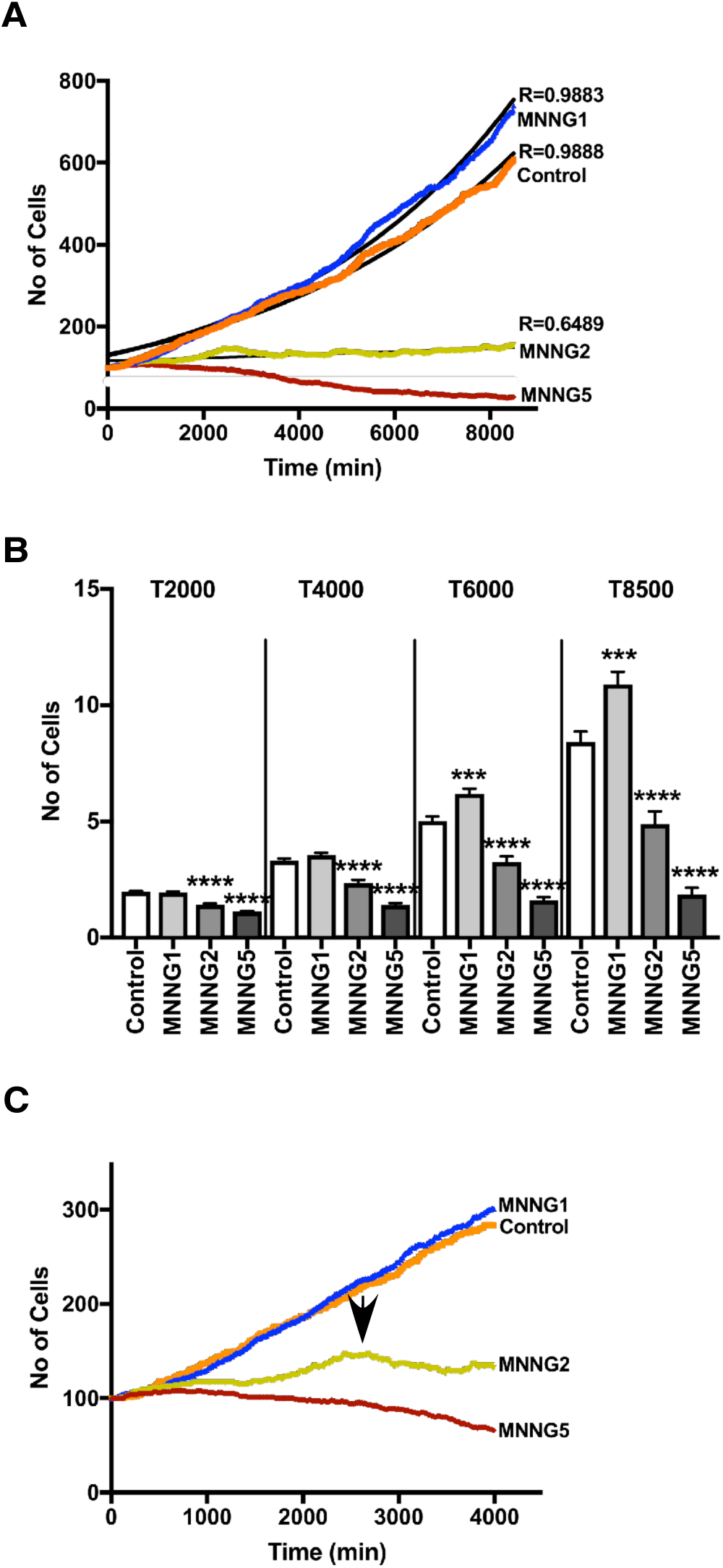
Effect of MNNG exposure on cell proliferation. **A**. The number of cells at each time point was determined using the cell-lineage database. The initial number of cells was normalized by 100. Curves were fitted using non-linear regression curve fitting (Prism 7). **B**. The numbers of cells produced from each progenitor cell at the 2,000, 4,000, 6,000, and 8,500 min time points were calculated and analyzed by Student’s *t*-tests; ***p<0.001 and ****p<0.0001 in relation to Control. Results shown as mean ± SEM. **C**. Growth curves for the time points from 1–4,000 min are shown. Arrow indicates the time point at which the number of cells exposed to MNNG2 reached the maximum.

### Characterization of cells exposed to MNNG5 using spatiotemporal data

We showed that a cytotoxic dose of MNNG (MNNG5) prolonged cell doubling time (Fig. 1B), inhibited cell population expansion (Fig. 2A), and induced CD (Fig. 3A). To clarify the individual cell- or cell population-based context leading to the induction of CD, we analyzed the spatiotemporal information on the cellular events experienced by individual cells. The results shown in Fig. 3 were normalized to the total number of cells to evaluate the chances of CD, MD, and CF events in individual cells. The results suggest that cytotoxic doses of MNNG induced MD (Fig. 3B, MNNG5) and CF (Fig. 3C, MNNG5). Then, we investigated the relationships between MD and CF events and CD by identifying the cellular events occurring prior to CD. MD and CF occurred prior to the induction of CD (Fig. 3D, MD > CD and CF > CD, MNNG5). Furthermore, 47.4% and 21.0% of CD events occurred following entry of MNNG5-exposed cells into mitosis (MI) and bipolar cell division (BD), respectively (Table 1, MNNG5, BD (exc. MI) > CD and MI > CD). Analysis of the responses of cells to MNNG5 thus revealed that CD induced by cytotoxic doses of MMNG occurred following various cellular events, implying that the determination of CD using conventional end point analysis may be biased towards CD induced following certain cellular events, given that conventional end point analyses are unable to distinguish among the different processes.

**Table 1.** Summary of cellular responses induced by MNNG.

| Treatments | Cell-doubling time |  | Rate of cell proliferation (fold increase) | CD |  |  |
| --- | --- | --- | --- | --- | --- | --- |
|  | min | fold (relative to Control) |  | Processes | % | No. of events/100 lineages (total no. of cells/100 lineages) |
| MNNG40 | nd<sup>1)</sup> | nd<sup>1)</sup> | nd<sup>1)</sup> | Movie S5 |  |  |
| MNNG5 | 2493 | 1.10 | 0.6 | BD (exc. MI) > CD<sup>2)</sup> | 21.0 | 25.5 |
|  |  |  |  | **Total BD (exc. MI) > CD** | **21.0** | **25.5** |
|  |  |  |  | BD (MI) > CD<sup>3)</sup> | 8.7 | 10.5 |
|  |  |  |  | No Div. MI > CD<sup>4)</sup> | 38.7 | 47.0 |
|  |  |  |  | **Total MI > CD** | **47.4** | **57.5** |
|  |  |  |  | MD > CD<sup>5)</sup> | 10.3 | 12.5 |
|  |  |  |  | CF > CD | 8.2 | 10.0 |
|  |  |  |  | No Div. > CD<sup>6)</sup> | 11.9 | 14.5 |
|  |  |  |  | Others > CD<sup>7)</sup> | 1.2 | 1.5 |
|  |  |  |  | **Total** | **100** | **121.5 (234.0)<sup>8)</sup>** |
| MNNG2 | 2186 | 0.96 | 1.5 | BD (DV1) > CD (G1) | 12.3 | 22.3 |
|  |  |  |  | BD (DV1) > CD (S+)<sup>9)</sup> | 9.0 | 16.4 |
|  |  |  |  | BD (DV2) > CD (G1) | 6.5 | 11.0 |
|  |  |  |  | BD (DV2) > CD (S+)<sup>9)</sup> | 6.5 | 11.0 |
|  |  |  |  | BD (DV3) > CD (G1) | 5.6 | 10.1 |
|  |  |  |  | BD (DV3) > CD (S+)<sup>9)</sup> | 7.6 | 13.8 |
|  |  |  |  | BD (DV4) > CD (G1) | 3.8 | 6.9 |
|  |  |  |  | BD (DV4) > CD (S+)<sup>9)</sup> | 1.2 | 2.1 |
|  |  |  |  | **Total BD > CD** | **52.5** | **93.6** |
|  |  |  |  | No Div. MI > CD<sup>4)</sup> | 12.9 | 23.4 |
|  |  |  |  | MD > CD<sup>5)</sup> | 16.1 | 29.3 |
|  |  |  |  | CF > CD | 13.8 | 25.0 |
|  |  |  |  | No Div. > CD<sup>6)</sup> | 4.7 | 8.5 |
|  |  |  |  | Others > CD<sup>7)</sup> | 0 | 0 |
|  |  |  |  | **Total** | **100** | **179.8 (671.2)<sup>8)</sup>** |
| MNNG1 | 2059 | 0.90 | 7.3 | BD (DV1) > CD (G1) | 2.6 | 6.0 |
|  |  |  |  | BD (DV1) > CD (S+)<sup>9)</sup> | 6.9 | 16.1 |
|  |  |  |  | BD (DV2) > CD (G1) | 5.1 | 12.0 |
|  |  |  |  | BD (DV2) > CD (S+)<sup>9)</sup> | 11.3 | 26.5 |
|  |  |  |  | BD (DV3) > CD (G1) | 6.9 | 16.1 |
|  |  |  |  | BD (DV3) > CD (S+)<sup>9)</sup> | 16.3 | 38.2 |
|  |  |  |  | BD (DV4) > CD (G1) | 11.0 | 25.9 |
|  |  |  |  | BD (DV4) > CD (S+)<sup>9)</sup> | 8.3 | 19.6 |
|  |  |  |  | **Total BD > CD** | **68.4** | **160.4** |
|  |  |  |  | No Div. MI > CD<sup>4)</sup> | 2.4 | 5.7 |
|  |  |  |  | MD > CD<sup>5)</sup> | 17.2 | 40.4 |
|  |  |  |  | CF > CD | 9.8 | 23.2 |
|  |  |  |  | No Div. > CD<sup>6)</sup> | 1.8 | 4.1 |
|  |  |  |  | Others > CD<sup>7)</sup> | 0.4 | 0.9 |
|  |  |  |  | **Total** | **100** | **232.7**<br>**(1907.0)<sup>8)</sup>** |
| Control | 2276 | 1.0 | 6.0 | BD (DV1) > CD (G1) | 3.4 | 6.9 |
|  |  |  |  | BD (DV1) > CD (S+)<sup>9)</sup> | 11.3 | 23.1 |
|  |  |  |  | BD (DV2) > CD (G1) | 5.4 | 11.1 |
|  |  |  |  | BD (DV2) > CD (S+)<sup>9)</sup> | 15.9 | 32.4 |
|  |  |  |  | BD (DV3) > CD (G1) | 9.8 | 19.9 |
|  |  |  |  | BD (DV3) > CD (S+)<sup>9)</sup> | 16.3 | 33.3 |
|  |  |  |  | BD (DV4) > CD (G1) | 9.5 | 19.4 |
|  |  |  |  | BD (DV4) > CD (S+)<sup>9)</sup> | 4.8 | 9.7 |
|  |  |  |  | **Total BD > CD** | **76.4** | **155.8** |
|  |  |  |  | No Div. MI > CD<sup>4)</sup> | 1.0 | 1.9 |
|  |  |  |  | MD > CD<sup>5)</sup> | 10.2 | 20.8 |
|  |  |  |  | CF > CD | 11.1 | 22.7 |
|  |  |  |  | No Div. > CD<sup>6)</sup> | 0.9 | 1.9 |
|  |  |  |  | Others > CD<sup>7)</sup> | 0.5 | 0.9 |
|  |  |  |  | **Total** | **100** | **204.0**<br>**(1611.1)<sup>8)</sup>** |

<sup>1)</sup> nd: not determined.
<sup>2)</sup> BD (exc. MI) > CD: CD occurred either following BD (prior to MI) or following BD (after entering into MI). BD > CD includes CD occurred prior to MI.
<sup>3)</sup> BD (MI) > CD: CD occurred either following BD (prior to MI) or following BD (after entering into MI). BD (MI) > CD includes CD occurred following MI.
<sup>4)</sup> No Div. MI > CD includes CD occurred following MI, but preceding events of MI were unable to be determined.
<sup>5)</sup> MD > CD includes CD occurred following MD.
<sup>6)</sup> No Div. > CD includes CD, of which preceding events were unable to be determined.
<sup>7)</sup> Others > CD includes CD occurred following incomplete mitosis.
<sup>8)</sup> Total number of cells per 100 cell lineages are shown.
<sup>9)</sup> S+ includes CD occurred during S+G2+M.

**Fig. 3.**
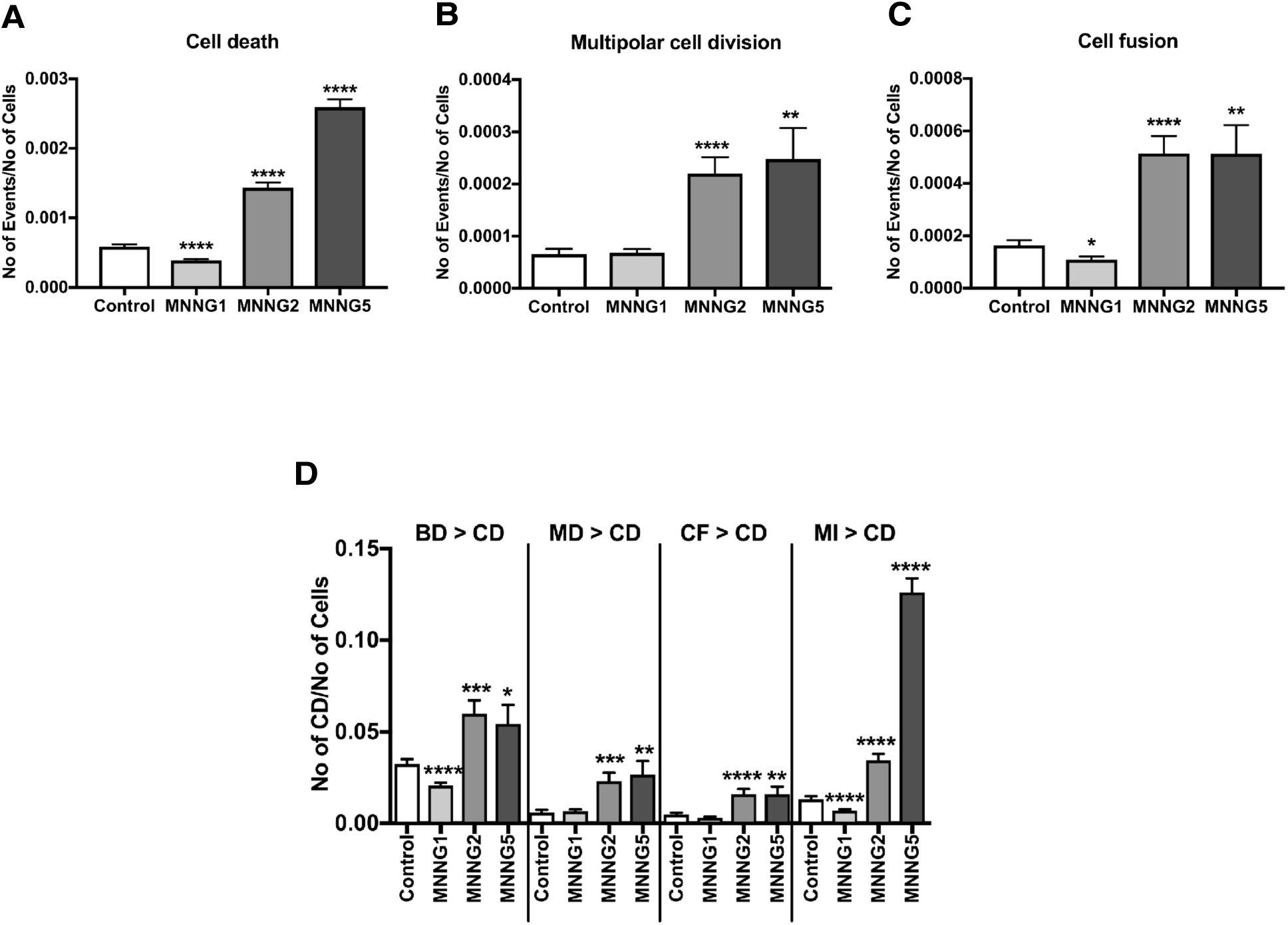
Numbers of CD, MD, and CF in cells exposed to MNNG. Data in **A-C** were normalized by the total number of cells. The numbers of CD (**A**), MD (**B**), and CF (**C**) events were determined using the cell lineage database. **D**. The numbers of CD events that occurred following BD, MD, CF, or MI are shown. If the event preceding MI was MD or CF, the CD event following the MI was included in MD > CD or CF > CD, respectively. **A-D**. Student’s *t*-tests were performed in relation to Control; *p<0.05, **p<0.01, ***p<0.001 and ****p<0.0001. Results shown as mean ± SEM.

### Characterization of cells exposed to MNNG2 using spatiotemporal data

In contrast to cells exposed to MNNG5, cells treated with MNNG2 were able to proliferate (Fig. 2C), and did not appear to undergo immediate CD. Clinically relevant doses of alkylating agents do not induce immediate responses (35, 36), suggesting that MNNG2 may represent a clinically relevant dose equivalent. We therefore analyzed the responses of HeLa cells exposed to MNNG2 using spatiotemporal data. Similar to cells exposed to MNNG5, the numbers of CD, MD, and CF events were significantly increased by exposure to MNNG2 (Fig. 3A-C, MNNG2). However, in contrast to MNNG5-exposed cells, CD occurred more frequently in cells following BD, compared with following entry into MI (Fig. 3D). We further explored the occurrence of CD induced by MNNG2 by comparing the frequencies of BD and CD at each time point, given that the growth of MNNG2 cells is likely to represent a balance between cell generation and CD (Fig. 2A). The numbers of BD and CD events plotted at each time point are shown in Fig. 4A-D. BD occurred constantly throughout the observation period in Control cells (Fig. 4A) and CD started to occur after approximately 2,000 min (Fig. 4B). BD also occurred throughout the observation period in MNNG2-exposed cells, but there were fewer BD cells compared with control cells (Fig. 4A and 4C), while CD also started to occur after approximately 2,000 min (Fig. 4D). We evaluated the relationship between BD and CD quantitatively by calculating the CD/BD ratio. The CD/BD ratio in control cells before 2,000 min was 7.4/92.5 = 0.08, implying that BD was predominant. This ratio was increased to 10.1/37.2 = 0.27 by exposure to MNNG2, due to a reduction in BD. However, BD was still predominant, resulting in a small increase in cell population size (Fig. 2C). The CD/BD ratio in the control cells after 2,000 min was 197.2/625.0 = 0.32, reflecting the predominance of cell growth over CD, while the ratio in the MNNG2-exposed cells was 171.4/204.2 = 0.84, which was close to 1.0, indicating that proliferation of MNNG2-exposed cells was largely balanced by CD, resulting in only a 1.5-fold increase in cell population size (Fig. 2A).

**Fig. 4.**
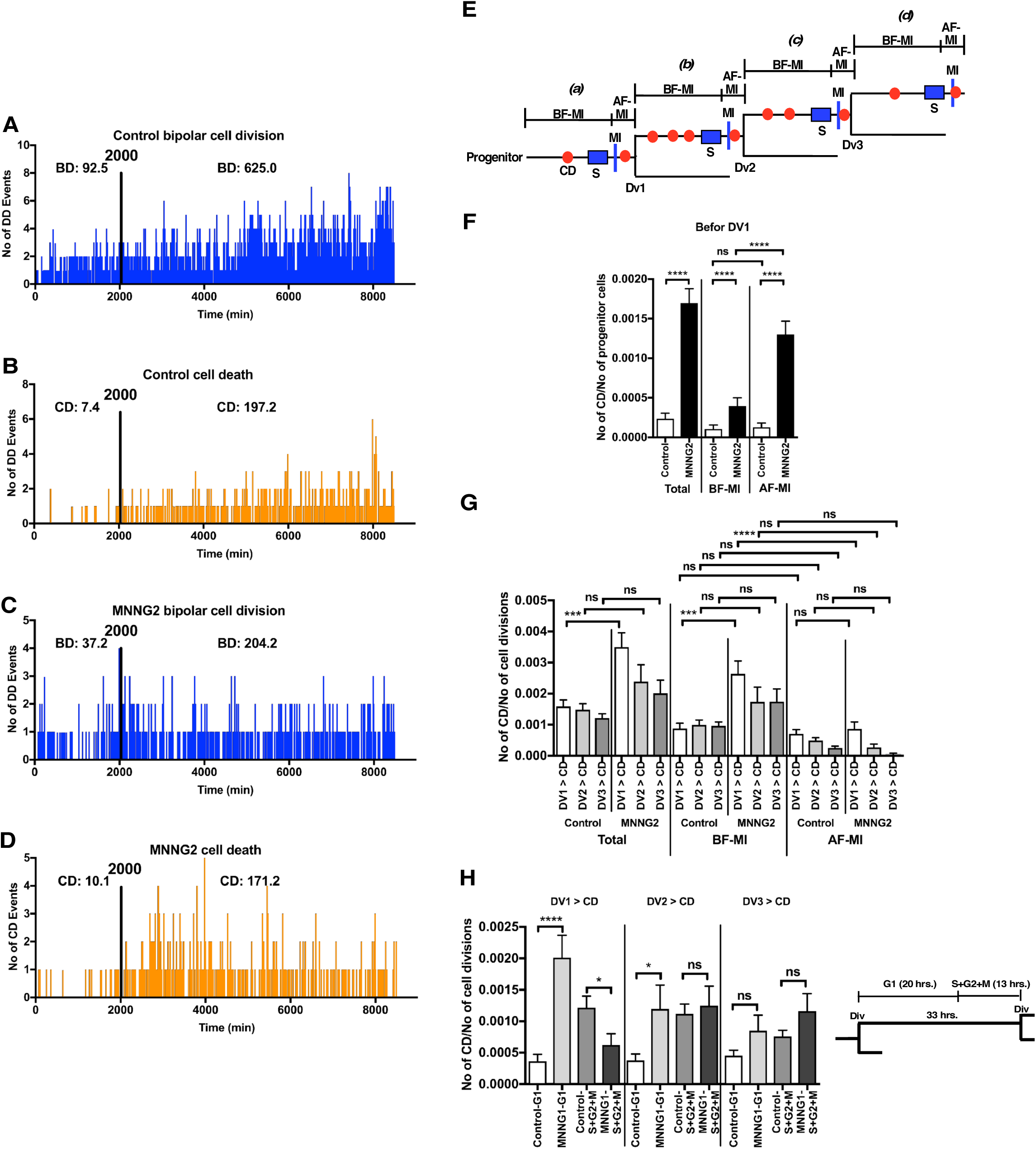
Analysis of cells exposed to MNNG2. The numbers of BD (**A and C**) and CD (**B and D**) events occurring at each time point in Control cells (**A and B**) and cells exposed to MNNG2 (**C and D**) are plotted. The total numbers (per 100 cell lineages) of BD and CD events that occurred before and after 2,000 min were determined (**A-D**). **e**. CD events (red circles) occurring in progenitor cells *(a)*, and following the division of progenitor cells (DV1, *(b)*), daughter cells (DV2, *(c)*), and GD cells (DV3, *(d))* were determined. Then, CD events occurred before (BF-MI) and after (AF-MI) entering MI were also determined for *a, b, c and d*. S represents S phase. **F**. The numbers of CD events that occurred in progenitor cells were normalized by the numbers of progenitor cells. Results shown as mean ± SEM. One-way ANOVA was performed; ns: not significant and ****p<0.0001. **G**. The numbers of CD events that occurred following DV1, DV2, and DV3 were normalized by the numbers of DV1, DV2, and DV3, respectively. Results shown as mean ± SEM. One-way ANOVA was performed; ns: not significant, ***p<0.001 and ***p<0.0001. **F and G** Total: the number of CD events occurred before and after entering into MI. **H**. The cell doubling time of cells exposed to MNNG2 was assumed to be 33 h; the duration of S1+G2+M phases was 13 h, and the duration of G1 was thus 20 h. CD events occurring during G1 and S+G2+M phases were determined. Results shown as mean ± SEM. Student’s *t*-tests were performed; ns: not significant, *p<0.05, ***p<0.001 and ****p<0.0001.

Given that the mean cell doubling time of MNNG2-exposed cells was about 2,000 min (Fig. 1B and e), these results also suggest that CD was likely to occur after a cell division. Indeed, previous end point analyses suggest that CD induced by exposure to low levels of alkylating agents occurred after the first S phase by recognition of *O*^6^-methyl G:T and *O*^6^-methyl G:C mismatches (22, 23), the second S phase by removal of mismatched by mismatch correction (20, 21, 31) or the third S phase entering after the formation of multinuclear cell (28). We therefore determined if CD occurred prior to DV1, or following DV1, DV2, or DV3 (Fig. 4E). Then, the numbers of CD events prior to DV1 were determined and normalized to the numbers of progenitor cells (Fig. 4E *(a))*. The numbers of CD events after DV1, DV2, and DV3 (Fig. 4E *(b), (c)*, and *(d))* were normalized to the numbers of divisions, i.e. DV1, DV2, and DV3, respectively, given that the number of cell divisions increased following cell growth. As shown in Fig. 4F (Total: the number of CD events occurred before and after entering into MI)), total numbers of CD occurred prior to DV1 were significantly increased in cells exposed to MNNG2, and the increase was also observed before and after entering the cells into MI (Fig. 4F, BF-MI and AF-MI; and Table 1, No Div. > CD and No Div. MI > CD), although CD events were more frequently occurred after entering into MI. These results support a hypothesis that CD induced by exposure to low levels of alkylating agents occurred after the first S phase (22, 23). We then determined the number of CD events occurred following DV1, DV2 and DV3 in control HeLa cells and found that CD occurred spontaneously with a constant probability during cell culture (Fig. 4G, Total, Control). In contrast, in MNNG2 treated cells, CD events following DV1 were significantly increased compared with the control cells (Fig. 4G, Total, DV1 > CD, Control vs. MNNG2). There was no significant difference in CD events following DV2 or DV3, but CD tended to be increased following these divisions (Fig. 4G, Total). We also found similar tendencies in cells, which were before and after entering into MI (Fig. 4G, BF-MI and AF-MI). These results suggest that CD mainly occurred in MNNG2-exposed cells after DV1 as reported (20, 21, 31), but less frequently occurred after DV2 and DV3, and after entering into MI. With this regard, it has been proposed that CD occurred following G2 arrest (20, 21, 31), while it also has been reported that the arrest is not involved in CD induced by low dose of alkylating agents (28). To clarify these processes, we determined if CD occurred during S and G2 phases, as suggested previously. The mean cell doubling time of cells exposed to MNNG2 was 2,186 min (33 h); we assumed that the duration of the S1+G2+M phase was 13 h, and the duration of G1 phase was thus 20 h (Fig. 4H, right). We then calculated the numbers of CD events during the different phases. CD in control cells occurred mainly in S+G2+M phase (Fig. 4H). Although we also found that CD occurred in MNNG2-treated cells during S+G2+M phase, MNNG2-induced CD occurred predominantly during G1 (Fig. 4H), supporting a hypothesis that some CD events are induced without arresting cells at G2 phase (28). In summary, CD events were induced by various processes in MNNG2 cells, as summarized in Table 1, implying that investigations of CD in MNNG2-exposed cells using conventional analysis may produce different conclusions, depending on which specific processes are detected.

### Characterization of cells exposed to MNNG1 using spatiotemporal data

We defined a non-cytotoxic dose of MNNG as a dose at which cell proliferation was not reduced and investigated the cellular alterations and responses induced by such a non-cytotoxic dose of MNNG. MNNG1 shortened the cell doubling time (Fig. 1A and B) and promoted cell proliferation (Fig. 2A), but suppressed the induction of CD and CF (Fig. 3A and C), suggesting that MNNG1 mainly promoted cell growth. We previously investigated the roles of individual HeLa and A549 cells in maintaining the cell population by synchronizing the cell cycle *in silico*, by grouping cells based on the number of progeny produced by a progenitor cell, and identified a well-growing sub-population using this approach (1, 3). We used a similar approach to characterize cells exposed to MNNG1 in the current study. The *in silico* cell cycle synchronization process normalized the first cell division as time point 1 (Fig. 5A), and the cell growth curves determined after synchronization are shown in Fig. 5B. We then determined the number of progeny generated by a single progenitor cell at 8,500 min using the synchronized data. As shown in Table 2 (Group A, 0–2; B, 3–5; C, 6–8; D, 9–11; E, 12–14; F, 15–17; and G, ≥ 18 cells), 4.65% of control cells produced ≥18 progeny cells (Group G), which was used as an indicator of high reproductive ability. In contrast, 11.68% of cells exposed to MNNG1 produced ≥18 progeny cells (Group G), suggesting that MNNG1 exposure increased the number of cells with high reproductive ability. Cells with a high reproductive ability had a significantly shorter cell doubling time than control cells (Table 2 and Fig. 5C, e.g., Group G cells). MNNG1 was therefore likely to stimulate the growth of a certain sub-population of cells by increasing their reproductive ability and shortening their cell doubling time. We previously determined if the progeny retained the reproductive ability of their progenitor cell by comparing the reproductive abilities of the progenitor and granddaughter (GD) cells (1). In the current study, we first determined the number of progeny produced by a progenitor cell at 3,500 min (Fig. 5A), and then identified the GD cells of that progenitor cell and counted the number of progeny produced by the GD cells after 3,500 min from the first cell division of the GD. If a GD cell retained a similar reproductive ability to its progenitor cell, the GD cell would be expected to produce a similar number of progeny to its progenitor cell. Accordingly, we found no significant difference between the numbers of progeny produced by progenitor and DG cells (Fig. 6A, progenitor vs. DG), indicating that DG cells retained a similar reproductive ability to their progenitor cells. However, progenitor and GD cells exposed to MNNG1 produced more progeny than the respective control cells, suggesting that MNNG1 promoted proliferation of both progenitor and DG cells. We reorganized the data in Fig. 6A according to the number of progeny produced by a progenitor cell (Fig. 6B), and showed that MNNG1 increased the number of progenitor cells that were able to produce more than five progeny (Fig. 6B, progenitor, 7.8 % for Control and 31.7% for MNNG1). We previously reported that HeLa cells contained putative cancer stem cells, accounting for 3%–7% of the cell population, which had high reproductive ability (produced more than seven progeny cells) and were involved in maintaining the HeLa cell population (1). The percentages of progenitor and GD cells that produced seven or eight progeny are shown in Fig. 6B. These cells accounted for 3.6%–4.6% of control cells compared with 9.5%–15.1% of cells exposed to MNNG1. The additional cells (5.9%–10.5%) were likely to be cells with lower reproductive capacity that were converted to more highly reproductive cells by exposure to MNNG1. These results suggest that the increased growth of cells exposed to MNNG1 is the result of the generation of highly reproductive cells, referred to as putative cancer stem cells (1).

**Table 2.** The number of progeny produced from a progenitor cell and the mean cell-doubling times.

| Group:<br>No. of<br>progeny<sup>1)</sup> | Control |  | MNNG1 |  | MNNG2 |  | MNNG5 |  |
| --- | --- | --- | --- | --- | --- | --- | --- | --- |
|  | % <sup>2)</sup> | Doubling<br>time (h)<sup>3)</sup> | %<sup>2)</sup> | Doubling<br>time (h)<sup>3)</sup> | %<sup>2)</sup> | Doubling<br>time (h)<sup>3)</sup> | %<sup>2)</sup> | Doubling<br>time (h)<sup>3)</sup> |
| A: 0–2 | 37.50 | **40.1 ± 0.9** | 39.42 | **34.7 ± 0.5\*\*\*\*** | 80.31 | **35.6 ± 0.7\*\*\*\*** | 96.50 | **43.2 ± 2.5** |
| B: 3–5 | 17.12 | **39.6 ± 0.6** | 12.93 | **38.9 ± 0.6** | 9.55 | **36.9 ± 0.7\*\*** | 2.50 | **41.6 ± 1.9** |
| C: 6–8 | 14.35 | **37.3 ± 0.4** | 9.15 | **37.7 ± 0.5\*** | 3.72 | **36.5 ± 1.0\*** | 1.00 | **34.6 ± 1.5\*** |
| D: 9–11 | 11.57 | **38.1 ± 0.4** | 11.36 | **35.9 ± 0.3\*** | 3.19 | **37.5 ± 0.9** | 0 | **0** |
| E: 12–14 | 8.33 | **37.3 ± 0.4** | 7.89 | **34.4 ± 0.3\*\*\*\*** | 2.17 | **33.8 ± 0.7\*\*\*\*** | 0 | **0** |
| F: 15–17 | 6.48 | **36.6 ± 0.4** | 7.57 | **33.9 ± 0.3\*\*\*\*** | 0.53 | **33.4 ± 1.0\*** | 0 | **0** |
| G: ≥ 18 | 4.65 | **33.0 ± 0.4** | 11.68 | **31.2 ± 0.2\*\*\*** | 0.53 | **36.9 ± 1.2\*\*\*** | 0 | **0** |
| Total | 100 |  | 100 |  | 100 |  | 100 |  |

<sup>1)</sup> Progenitor cells were grouped (Group A-G) by the number of progeny produced at time 8500 min.
<sup>2)</sup> The % of progenitor cells that produced the number of progeny shown in the “Group: No. of progeny” column is shown.
<sup>3)</sup> The mean cell-doubling times and SEMs were calculated using the cell-doubling times of each cell.
Student’s *t*-tests were performed; \* $p < 0.05$ , \*\* $p < 0.01$ , \*\*\* $p < 0.001$ and \*\*\*\* $p < 0.0001$ in relation to Control.

**Fig. 5.**
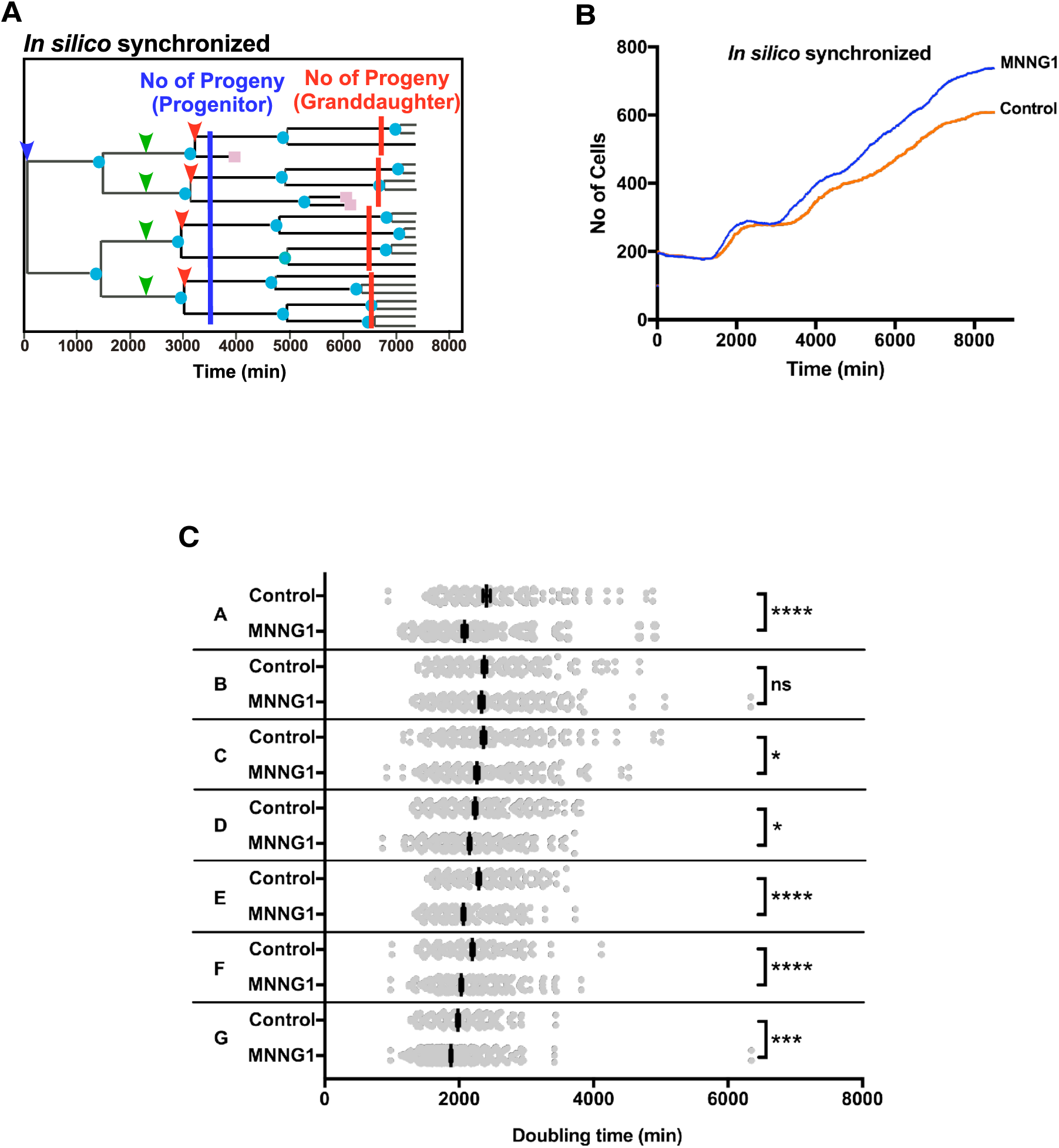
Analysis of progeny produced from Control progenitor and MNNG1-exposed cells. **A**. The cell cycle was synchronized *in silico*. The time point when the progenitor cells divided was normalized as Time 1 (blue arrowhead). Blue line indicates the time point 3500 min after the division. GD cells are indicated by green arrowheads. The red line shows the time point 3500 min after the division of GD cells (red arrowheads). **B**. Cell growth curves determined after synchronization. **C**. Progenitor cells were grouped (A-G) according to the number of progeny cells, as shown in Table 2. The cell-doubling times of each group of cells were then determined to perform Student’s *t*-tests; ns: not significant, and *p<0.05, ***p<0.001 and ****p<0.0001 in relation to Control. Results shown as mean ± SEM.

**Fig. 6.**
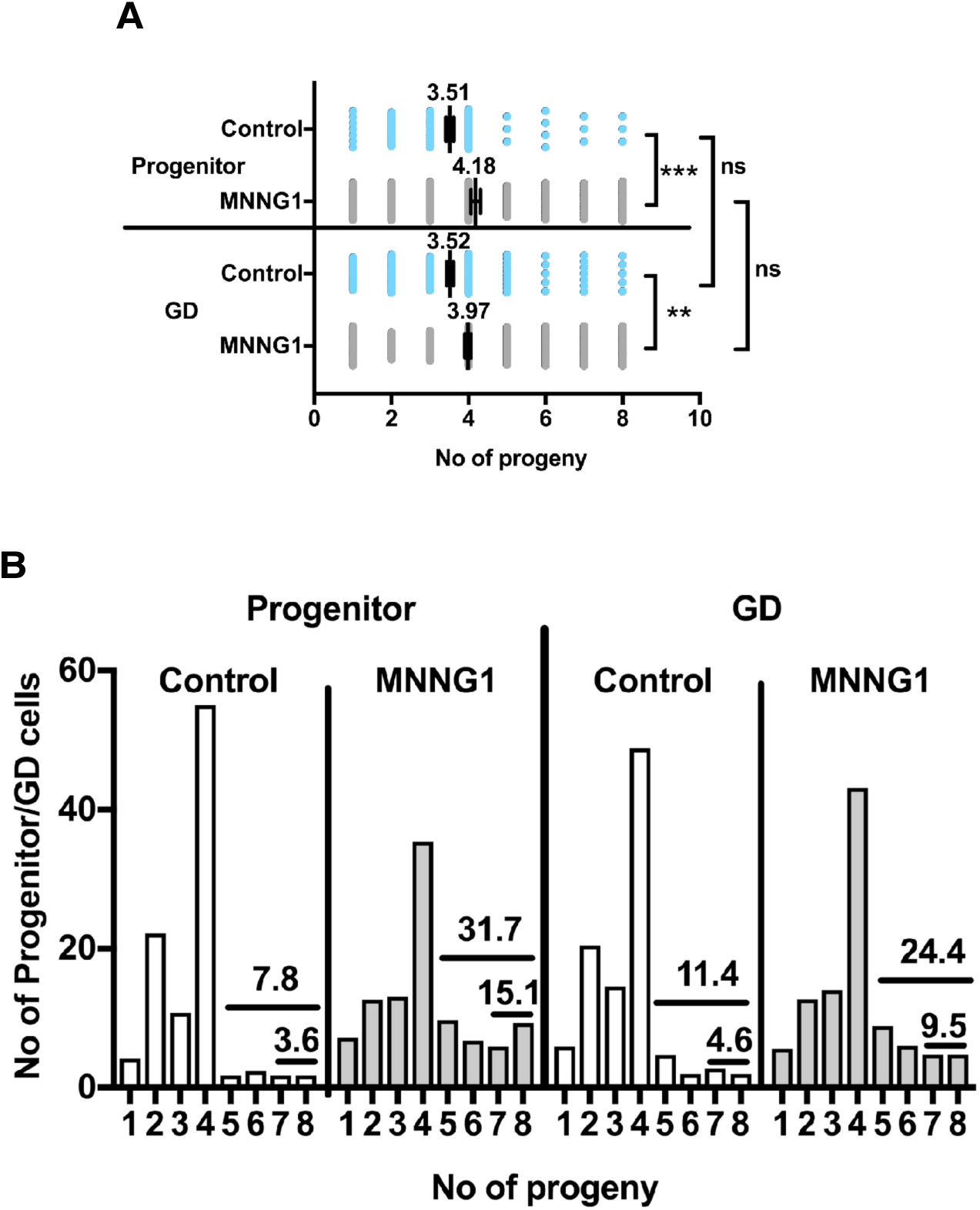
Analysis of the reproductive ability of Control and MNNG1-exposed cells. An *in silico* synchronized-cell lineage database was used. **A and B**. The number of progeny produced from a progenitor at 3500 mins and from DG cells at 3500 min from their cell division was determined. **A**. One-way ANOVA was performed; ns: not significant, **p<0.01 and ***p<0.01. Results shown as mean ± SEM. **B**. Percentages of progenitor and GD cells that produced 5–8 and 7–8 progeny are shown.

## Discussion

Various end point analyses have been used to investigate cellular responses to exogenously added substances. These analyses generate data on the characteristics of cells at a specific moment in time, and the events are thus deduced based on data obtained at the specific time when the cells were analyzed. In addition, these deductions are often based on the assumption that cells respond to an exogenously added substance in a stoichiometric manner; however cells may not respond in such a manner, given that cultured cells are composed of heterogeneous cell sub-populations with intrinsically different sensitivities to a substance. End point analyses may thus have a limited ability to characterize the responses of such cells to an exogenous substance. In contrast, single-cell tracking analysis can provide spatiotemporal data, which allows the responses of cells to be analyzed without the need for relying on deductions and assumptions. However, such an approach for characterizing cells exposed to mutagenic and carcinogenic substances has not yet been developed. In the present study, we validated this approach by characterizing cells exposed to various doses of MNNG using spatiotemporal data obtained from single-cell lineage tracking analysis. CD induced by different doses of MNNG is summarized in Table 1. Lethal doses of MNNG (40 μM) induced CD within 4 h (Movies S5).

Cytotoxic doses of MNNG predominantly induced CD following MI (BD (MI) > CD+ No Div. MI > CD = 47.4%) or BD (BD (exc. MI) > CD, 21.0%). Thus, an end point analysis to detect CD in cells exposed to cytotoxic doses of MNNG would have about 50% and 20% chances of detecting CD events occurring following MI and BD events, respectively, making it difficult for conventional end point analyses to determine which type of CD was analyzed. In the case of cells exposed to sub-cytotoxic doses of MNNG (MNNG2), CD events were induced via more diverse processes than those induced by MNNG5. Although CD occurring after BD (DV1, G1; i.e., CD occurs at G1 phase after the first BD) was a major cause of CD induced by MNNG2, CD after MD and CF were also induced at similar frequencies. End point analyses of cells exposed to MNNG2 may thus reach different conclusions depending on which CD process is analyzed. Indeed, previous reports suggest that CD occurred in cells exposed to low doses of an alkylating agent during S or G2 phase of daughter cells produced by BD of exposed progenitors, or following DV1, or DV2 (20–23, 25–31). It is likely that these reports investigated a specific CD process among the various processes identified here. We therefore consider that spatiotemporal data are essential for studying cellular responses to sub-cytotoxic doses of MNNG.

Finally, we demonstrated that non-cytotoxic doses of MNNG stimulated cell proliferation by promoting the growth of a sub-population of cells referred to as putative cancer stem cells (1). MNNG is known to induce hyperplasia in rodents (37, 38), and non-cytotoxic dose of MNNG may thus act as a cell growth promoter. However, the doses of carcinogens used in laboratory tests are generally 100–1,000 times higher than the doses present in the environment (39), as environmental doses are unlikely to induce detectable responses in cultured cells. The biological responses induced by environmental doses of carcinogens are thus poorly understood. Although MNNG is not an environmental carcinogen, our results suggest that cellular responses induced by doses that are too low to cause significant induction of CD can still be analyzed based on spatiotemporal data for individual cells.

In conclusion, we propose that a single-cell lineage tracking analysis system that creates spatiotemporal data for individual cells represents a novel and potentially valuable approach for elucidating the effects of cytotoxic and non-cytotoxic doses of various carcinogenic and mutagenic substances. For example, CD is known to be induced by various mechanisms (40), and taking account of the events occurring prior to the induction of CD will allow a deeper understanding of CD overall. Although single-cell lineage tracking analysis remains a tedious process, computerization of the system may allow it to become a routine method for characterizing cells exposed to genotoxic substances.

## Materials and Methods

HeLa cells were cultured in DMEM containing 10% fetal bovine serum in a humidified 5% CO_2_. To plate cells onto a coverglass Lab-Tek 8 well chamber, 50 μl of HeLa suspension containing 3500 cells were placed at the center of each well and left until cells attached to the coverglass surface. Then, 0.75 ml of culture medium was added to each well. Cells were used for live cell imaging 18 h after the plating. HeLa cells were treated with various concentration of MNNG (Sigma-Aldrich) for 30 min in serum-free DMEM.

## Long-term live cell imaging and data analysis

Quorum WaveFX Spinning Disc Confocal System (Quorum Technologies Inc., Canada) with Leica microscope controlled by Volocity v4.0 was used for long-term live cell imaging. DIC images were taken through HCX PL APO 40x oil objectives (NA=1.25) by using a Halogen lamp as a light source. Cells that were grown on a coverglass Lab-Tek 8 well chamber were then placed on a microscope stage and were cultured using an environmental chamber at 37°C with 7.5% humidified CO_2_ (Pathology Devices Inc, MD). In each well, a two-dimensional image acquisition array (filed of views: FOVs) was made to cover the area of interest (1). XY positions of FOVs were then registered using Volocity v4.0. DIC images were captured every 10 min from + 10 to − 10 μm of the focal plane at every 1 μm using piezo focus drive. Exposure time was 34 msec. To make time-lapse movies, focused images were selected from 21 z-plane image files using Volocity v4.0. After the selection, files containing focused image were assembled into a movie using QuickTime Player Pro. Panorama views of time point = 1 were prepared and cell lineage numbers were assigned to cells in a selected area (1). After assigning the cell lineage numbers, each cell was tracked using QuickTime Player Pro and the time points that MI, BD, MD, CF, and CD occurred in each cell were determined. To draw cell lineage maps and process data, C/C++ programs were written. Live cell imaging was started at 80% of confluency and we have previously confirmed that cell growth was continued for at least 9000 min (see Movie S1–2, Fig. 2 and ref. (1)). About 200–300 each of progenitor cells were tracked.

## Determination of cell-doubling times

We used cell-lineage database to determine cell-doubling time of individual cells. The time when a cell was produced by cell division (Time A) and the time when the same cell produced daughter cells (Time B) were determined, and cell-doubling time was calculated by subtracting Time A from Time B.

## Statistical analyses of cell-doubling times

Cell-doubling times were analyzed by Student’s *t*-tests (unpaired and two-tailed) or one-way ANOVA using Prism 7.

## Statistical analyses of cell growth

The number of progenitor cells and/or progeny of each cell lineage found at 2000, 4000, 6000 and 8500 min was determined. The data was analyzed by Student’s *t*-tests (unpaired and two-tailed) using Prism 7.

## Statistical analyses of cellular events

The number of BD, MD, CD, and CF occurred in each cell lineage was determined. The data was analyzed by Student’s *t*-tests (unpaired and two-tailed) or one-way ANOVA using Prism 7.

## Acknowledgements

We like to acknowledge Ms. Julie-Christine Lévesque, and the Bioimaging Platform of Research Center for Infectious Diseases, CRCHU de Québec for the technical support of microscopes. This work was supported by the Canadian Foundation for Innovation for MSS and SS.

## Author contributions

M.S.S. and S.S. designed this work, performed the experiments and wrote the paper. A.R. performed experiments and single-cell lineage tracking analysis.

## Competing Fanatical Interests

The authors declare no competing financial interests.

## References

1. Sato S, Rancourt A, Sato Y, & Satoh MS (2016) Single-cell lineage tracking analysis reveals that an established cell line comprises putative cancer stem cells and their heterogeneous progeny. Sci Rep 6: 23328.

2. Guo S, et al. (2014) Nonstochastic reprogramming from a privileged somatic cell state. Cell 156(4):649–662.

3. Rancourt A, Sato S, & Satoh MS (Basal level of p53 regulates cell population homeostasis. *BioRxiv* doi: https://doi.org/10.1101/319525.

4. Drablos F, et al. (2004) Alkylation damage in DNA and RNA--repair mechanisms and medical significance. DNA Repair (Amst) 3(11): 1389–1407.

5. Kubota Y, et al. (1996) Reconstitution of DNA base excision-repair with purified human proteins: interaction between DNA polymerase beta and the XRCC1 protein. EMBO J 15(23):6662–6670.

6. Sedgwick B, Bates PA, Paik J, Jacobs SC, & Lindahl T (2007) Repair of alkylated DNA: recent advances. DNA Repair (Amst) 6(4):429–442.

7. Singer B, Fraenkel-Conrat H, Greenberg J, & Michelson AM (1968) Reaction of nitrosoguanidine (N-methyl-N'-nitro-N-nitrosoguanidine) with tobacco mosaic virus and its RNA. Science 160(3833):1235–1237.

8. Strauss B, Scudiero D, & Henderson E (1975) The nature of the alkylation lesion in mammalian cells. Basic Life Sci 5A: 13–24.

9. Burkle A (2001) Physiology and pathophysiology of poly(ADP-ribosyl)ation. Bioessays 23(9):795–806.

10. Lindahl T, Satoh MS, Poirier GG, & Klungland A (1995) Post-translational modification of poly(ADP-ribose) polymerase induced by DNA strand breaks. Trends Biochem Sci 20(10):405–411.

11. Luo X & Kraus WL (2012) On PAR with PARP: cellular stress signaling through poly(ADP-ribose) and PARP-1. Genes Dev 26(5):417–432.

12. Ray Chaudhuri A & Nussenzweig A (2017) The multifaceted roles of PARP1 in DNA repair and chromatin remodelling. Nat Rev Mol Cell Biol 18(10):610–621.

13. Satoh MS & Lindahl T (1992) Role of poly(ADP-ribose) formation in DNA repair. Nature 356(6367):356–358.

14. Yu SW, et al. (2002) Mediation of poly(ADP-ribose) polymerase-1-dependent cell death by apoptosis-inducing factor. Science 297(5579):259–263.

15. Eadie JS, Conrad M, Toorchen D, & Topal MD (1984) Mechanism of mutagenesis by O6-methylguanine. Nature 308(5955):201–203.

16. Myrnes B, Giercksky KE, & Krokan H (1982) Repair of O6-methyl-guanine residues in DNA takes place by a similar mechanism in extracts from HeLa cells, human liver, and rat liver. J Cell Biochem 20(4):381–392.

17. Duckett DR, Bronstein SM, Taya Y, & Modrich P (1999) hMutLalpha- and hMutLalpha-dependent phosphorylation of p53 in response to DNA methylator damage. Proc Natl Acad Sci U S A 96(22): 12384–12388.

18. Duckett DR, et al. (1996) Human MutSalpha recognizes damaged DNA base pairs containing O6-methylguanine, O4-methylthymine, or the cisplatin-d(GpG) adduct. Proc Natl Acad Sci U S A 93(13):6443–6447.

19. York SJ & Modrich P (2006) Mismatch repair-dependent iterative excision at irreparable O6-methylguanine lesions in human nuclear extracts. J Biol Chem 281(32):22674–22683.

20. Mojas N, Lopes M, & Jiricny J (2007) Mismatch repair-dependent processing of methylation damage gives rise to persistent single-stranded gaps in newly replicated DNA. Genes Dev 21(24):3342–3355.

21. Stojic L, et al. (2004) Mismatch repair-dependent G2 checkpoint induced by low doses of SN1 type methylating agents requires the ATR kinase. Genes Dev 18(11): 1331–1344.

22. Yang G, et al. (2004) Dominant effects of an Msh6 missense mutation on DNA repair and cancer susceptibility. Cancer Cell 6(2): 139–150.

23. Yoshioka K, Yoshioka Y, & Hsieh P (2006) ATR kinase activation mediated by MutSalpha and MutLalpha in response to cytotoxic O6-methylguanine adducts. Mol Cell 22(4):501–510.

24. Friedman HS, Kerby T, & Calvert H (2000) Temozolomide and treatment of malignant glioma. Clin Cancer Res 6(7):2585–2597.

25. Hawn MT, et al. (1995) Evidence for a connection between the mismatch repair system and the G2 cell cycle checkpoint. Cancer Res 55(17):3721–3725.

26. Iyer RR, Pluciennik A, Burdett V, & Modrich PL (2006) DNA mismatch repair: functions and mechanisms. Chem Rev 106(2):302–323.

27. Schroering AG, Edelbrock MA, Richards TJ, & Williams KJ (2007) The cell cycle and DNA mismatch repair. Exp Cell Res 313(2):292–304.

28. Schroering AG, et al. (2009) Prolonged cell cycle response of HeLa cells to low-level alkylation exposure. Cancer Res 69(15):6307–6314.

29. Tominaga Y, Tsuzuki T, Shiraishi A, Kawate H, & Sekiguchi M (1997) Alkylation-induced apoptosis of embryonic stem cells in which the gene for DNA-repair, methyltransferase, had been disrupted by gene targeting. Carcinogenesis 18(5):889–896.

30. Wang Y & Qin J (2003) MSH2 and ATR form a signaling module and regulate two branches of the damage response to DNA methylation. Proc Natl Acad Sci U S A 100(26): 15387–15392.

31. Zhukovskaya N, Branch P, Aquilina G, & Karran P (1994) DNA replication arrest and tolerance to DNA methylation damage. Carcinogenesis 15(10):2189–2194.

32. Ganem NJ, Godinho SA, & Pellman D (2009) A mechanism linking extra centrosomes to chromosomal instability. Nature 460(7252):278–282.

33. Shi Q & King RW (2005) Chromosome nondisjunction yields tetraploid rather than aneuploid cells in human cell lines. Nature 437(7061): 1038–1042.

34. Smith GJ & Grisham JW (1983) Cytotoxicity of monofunctional alkylating agents. Methyl methanesulfonate and methyl-N'-nitro-N-nitrosoguanidine have different mechanisms of toxicity for 10T1/2 cells. Mutat Res 111(3):405–417.

35. Jiricny J (2006) The multifaceted mismatch-repair system. Nat Rev Mol Cell Biol 7(5):335–346.

36. Kunkel TA & Erie DA (2005) DNA mismatch repair. Annu Rev Biochem 74: 681–710.

37. Chopra DP & Wilkoff LJ (1976) Inhibition and reversal by beta-retinoic acid of hyperplasia induced in cultured mouse prostate tissue by 3-methylcholanthrene or N-methyl-N'-nitro-N-nitrosoguanidine. J Natl Cancer Inst 56(3):583–589.

38. Chopra DP & Wilkoff LJ (1977) Reversal by vitamin A analogues (retinoids) of hyperplasia induced by N-methyl-N'-nitro-N-nitrosoguanidine in mouse prostate organ cultures. J Natl Cancer Inst 58(4):923–930.

39. Gold LS, Slone TH, Manley NB, & Ames BN (2002) Misconceptions About the Causes of Cancer (Fraser Institute, Vancouver, British Columbia, Canada).

40. Galluzzi L, et al. (2018) Molecular mechanisms of cell death: recommendations of the Nomenclature Committee on Cell Death 2018. Cell Death Differ 25(3):486–541.

